# Spasticity reduction in children with cerebral palsy is not associated with reduced energy consumption during walking

**DOI:** 10.1101/653071

**Authors:** Nicole L. Zaino, Katherine M. Steele, J. Maxwell Donelan, Michael H. Schwartz

## Abstract

**Background:** The average energy consumption during walking of children with cerebral palsy (CP) is over two times of that of typically developing (TD) children and fatigue is one of the top complaints of children with CP and their families. Spasticity has been theorized to contribute to increased energy consumption during walking in CP, but its role remains unclear.

**Methods:** We retrospectively compared the energy consumption of walking in children with diplegic CP before and after selective dorsal rhizotomy (SDR), a surgery that reduces spasticity. A control group of participants with CP who also underwent gait analysis but did not undergo SDR was matched to the SDR group by pre-surgery age, spasticity, and energy consumption. Energy consumption and spasticity were compared at baseline and follow-up for both groups.

**Findings:** As expected, the SDR group has a significantly greater decrease (−44%) in spasticity compared to matched peers with CP who did not undergo SDR (−16%, *P*<0.001). While both groups had a reduction in energy consumption between visits (12 % SDR and 14% no-SDR), there was no difference in the change in energy consumption between groups (*P*=0.4).

Interpretation: Reducing spasticity did not contribute to greater reductions in energy consumption, suggesting that spasticity has minimal impact on elevated energy consumption during walking for children with CP. Energy consumption and spasticity decrease with age among children with CP. Identifying matched control groups of peers with CP is critical for research involving children with CP to account for changes due to development.

**Highlights:**

- Energy consumption is not reduced after rhizotomy compared to matched peers
- Spasticity has minimal contribution to elevated energy during walking
- Matched control groups are critical in cerebral palsy research

## 1 Introduction

Cerebral palsy (CP) is a neuromuscular disorder caused by a brain injury at or near the time of birth, which primarily affects movement and coordination. CP is the most common pediatric disability in the United States, affecting over 2 per 1000 live births (Colver et al., 2014; Odding et al., 2006). Fatigue is one of the top complaints of children with CP and their families (Jahnsen et al., 2003). There are many different metrics used to evaluate energy during walking. Clinically, energy consumption is most often estimated from the volume of oxygen consumed per unit time and is an indicator of exertion (Schwartz et al., 2006). The energy consumed during walking for children with CP has been estimated to be two to three times that of typically developing (TD) peers (Bolster et al., 2017; Waters and Mulroy, 1999; Duffy et al., 1996; Norman et al., 2004; Rose et al., 1990). The cause of this increased energy consumption is unclear.

Spasticity, defined as a velocity-dependent resistance to stretch (Lance, 1980), has been theorized as a root cause of the observed increase in energy consumption in CP. Spasticity can cause an increase in overall muscle activity, that is thought to directly contribute to elevated energy consumption. Spasticity is observed in up to 80% of children with CP (Odding et al., 2006) and is also common among other neurologic disorders such as multiple sclerosis or spinal cord injury. Although the definition and presentation of spasticity differ, prior research has found that individuals with multiple sclerosis and spasticity also have energy costs two times that of TD peers, concluding that spasticity can be considered an important determinant of high energy costs of walking (Olgiati et al., 1988).

Selective dorsal rhizotomy (SDR) is a neurosurgical procedure where afferent nerve fibers in dorsal rootlets are cut to reduce efferent excitations (Tedroff et al., 2011). SDR has been shown to significantly reduce spasticity (Wang et al., 2018). Prior outcome studies have also suggested that SDR reduces energy consumption. For example, Carraro and colleagues found that energy was significantly reduced across multiple walking speeds after SDR for nine children with CP (Carraro et al., 2014). While these studies seem to indicate that spasticity contributes to elevated energy consumption in CP, there are several critical limitations. There are numerous other factors that could contribute to changes in energy consumption after surgery. For example, changes in walking patterns after SDR could change joint moments and muscle demands (Waters and Mulroy, 1999). Energy consumption is also known to decrease with age among children with CP (Kamp et al., 2014). Evaluating energy consumption before and after procedures thus require that these other factors are also considered. Identifying appropriate control groups of peers with CP provides one method to address these challenges. The purpose of this study was to determine if spasticity is a significant contributor to elevated energy consumption among children with CP by investigating if reducing spasticity leads to lower energy consumption. We hypothesized that if spasticity contributes to energy consumption for children with CP, then an SDR should result in greater changes in energy consumption compared to matched controls with CP. We matched individuals with CP who underwent gait analysis before and after SDR to peers with CP who did not undergo an SDR. Understanding the role of spasticity on energy consumption is important to inform treatments that aim to reduce fatigue and increase quality of life for children with CP.

## 2 Methods

### 2.1 Participants

We retrospectively identified individuals with diplegic CP who underwent gait analysis at Gillette Children’s Specialty Healthcare (St. Paul, MN, USA) between 1994 and 2018. Inclusion criteria for this research were:

1. primary diagnosis of diplegic CP,
2. underwent a bilateral SDR before the age of 12 years of age,
3. had at least one gait analysis before (baseline) and after (follow-up) SDR that included both oxygen consumption and Ashworth Scores,
4. had at least one gait analysis after SDR occurs before 18 years of age.

We also identified a control group of peers with CP that met the above inclusion criteria but did not undergo an SDR. This group was further matched to the treatment cohort by age, energy consumption, and spasticity at baseline. To identify matching peers between the control (no-SDR) and treatment (SDR) cohort, all matching variables for each participant were transformed into a single summary score using an autoencoder. An autoencoder is a neural network that can be used for dimensionality reduction of complex data (Møller, 1993). For this research, the autoencoder was defined using age, energy consumption, and spasticity across all children with diplegic CP who had previously received a gait analysis at Gillette to define a summary metric of the variations in these dimensions across the population. The summary scores were then calculated for each child in the SDR group and a k-nearest-neighbors (KNN) search algorithm was used to identify the closest matching peer with CP. If two children in the SDR group matched to the same peer, we checked to determine if there was another close match. A close match was defined as within 95% of the distance between all unique matches. All matching variables were compared between groups to evaluate the similarity of the cohorts.

### 2.2 Energy

Power during walking was assessed by converting breath-by-breath oxygen consumption 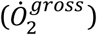 to Watts of power (*P*^*gross*^) using the conversion rate of 21 Joules/ml (Brockway, 1987). Both 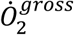 and 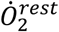 are converted to gas volume expressed under standard conditions of temperature, pressure, and dry (STPD) from testing conditions. The testing protocol consisted of a six minute walking trial preceded by a 3-10 minute rest period (Schwartz et al., 2006). Resting power (*P*^*rest*^) was assessed during supine or sitting (Schwartz et al., 2006). We calculated net-nondimensionalized (NN) power as:

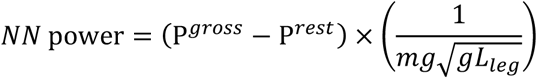

where *m* is body mass, *L*_*leg*_ is the length of the leg, and *g* is acceleration due to gravity. We evaluated nondimensionalized walking speed as (Hof, 1996):

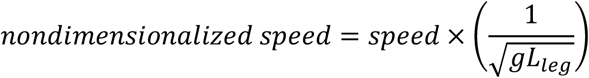

### 2.3 Spasticity

To evaluate spasticity, we calculated a summary spasticity score for each individual. The score was derived from the Ashworth Scale of six right limb muscle groups using principle component analysis (Rozumalski and Schwartz, 2009). The six muscle groups were hip adductors, hip flexors, hamstrings, rectus femoris, plantarflexors, and posterior tibialis. The Ashworth Scale is a discrete scale with five levels used to categorize spasticity (Pandyan et al., 1999). At Gillette, the following Ashworth Scale definitions are used: (1) no increase in tone, (2) slight increase in tone, (3) more marked increase in tone, (4) considerable increase in tone, and (5) affected part rigid. This spasticity summary score is a weighted average and can be interpreted as a typical Ashworth on a continuous 1-5 scale.

### 2.4 Statistical Analysis

A paired t-test (*α* < 0.05) was used to compare changes in energy consumption, spasticity, and walking speed between visits for each group (SDR and no-SDR). An independent samples t-test (*α* ≤ 0.05) was then used to compare change in energy consumption, spasticity, and walking speed between groups (SDR versus no-SDR). All values are reported as a mean (standard deviation) unless otherwise noted.

## 3 Results

### 3.1 Baseline Comparison

We identified 242 individuals with CP who met the inclusion criteria for the SDR group (136 male, 106 female, age: 6.0 (SD 1.7) years, height: 109.5 (SD 11.4) cm, weight: 19.3 (SD 6.0) kg) and matched 156 individuals with CP who did not undergo an SDR (Fig. 1, no-SDR group: 91 male, 65 female, age: 6.3 (SD 1.8) years, height: 112.1 (SD 11.9) cm, weight: 21.0 (SD 7.6) kg). The distributions for age, walking speed, energy consumption, and spasticity at baseline were similar between groups (Table 1).

**Table 1.** Summary of demographics and outcome measures for both cohorts.

|  | Baseline |  |  |  | Follow-up |  |  |  | Change |  |  |  |
| --- | --- | --- | --- | --- | --- | --- | --- | --- | --- | --- | --- | --- |
|  | SDR |  | no-SDR |  | SDR |  | no-SDR |  | SDR |  | no-SDR |  |
|  | ave | SD | ave | SD | ave | SD | ave | SD | ave | SD | ave | SD |
| Age (years) | 6.0 | 1.7 | 6.3 | 1.8 | 7.8 | 2.0 | 9.0 | 2.9 | 1.9 | 1.2 | 2.7 | 2.4 |
| Height (m) | 109.5 | 11.4 | 112.1 | 11.9 | 120.7 | 12.4 | 127.9 | 17.3 | 11.2 | 7.5 | 15.8 | 13.5 |
| Weight (kg) | 19.3 | 6.0 | 21.0 | 7.6 | 24.4 | 8.4 | 29.6 | 12.9 | 5.1 | 4.7 | 8.6 | 10.0 |
| Energy Consumption | 0.22 | 0.07 | 0.21 | 0.07 | 0.18 | 0.06 | 0.18 | 0.06 | -0.03 | 0.07 | -0.03 | 0.07 |
| Spasticity | 2.1 | 0.6 | 1.9 | 0.5 | 1.2 | 0.3 | 1.6 | 0.6 | -0.9 | 0.5 | -0.3 | 0.6 |
| Walking Speed | 0.28 | 0.10 | 0.30 | 0.09 | 0.30 | 0.10 | 0.30 | 0.08 | 0.02 | 0.09 | 0.002 | 0.07 |

**Figure 1.**
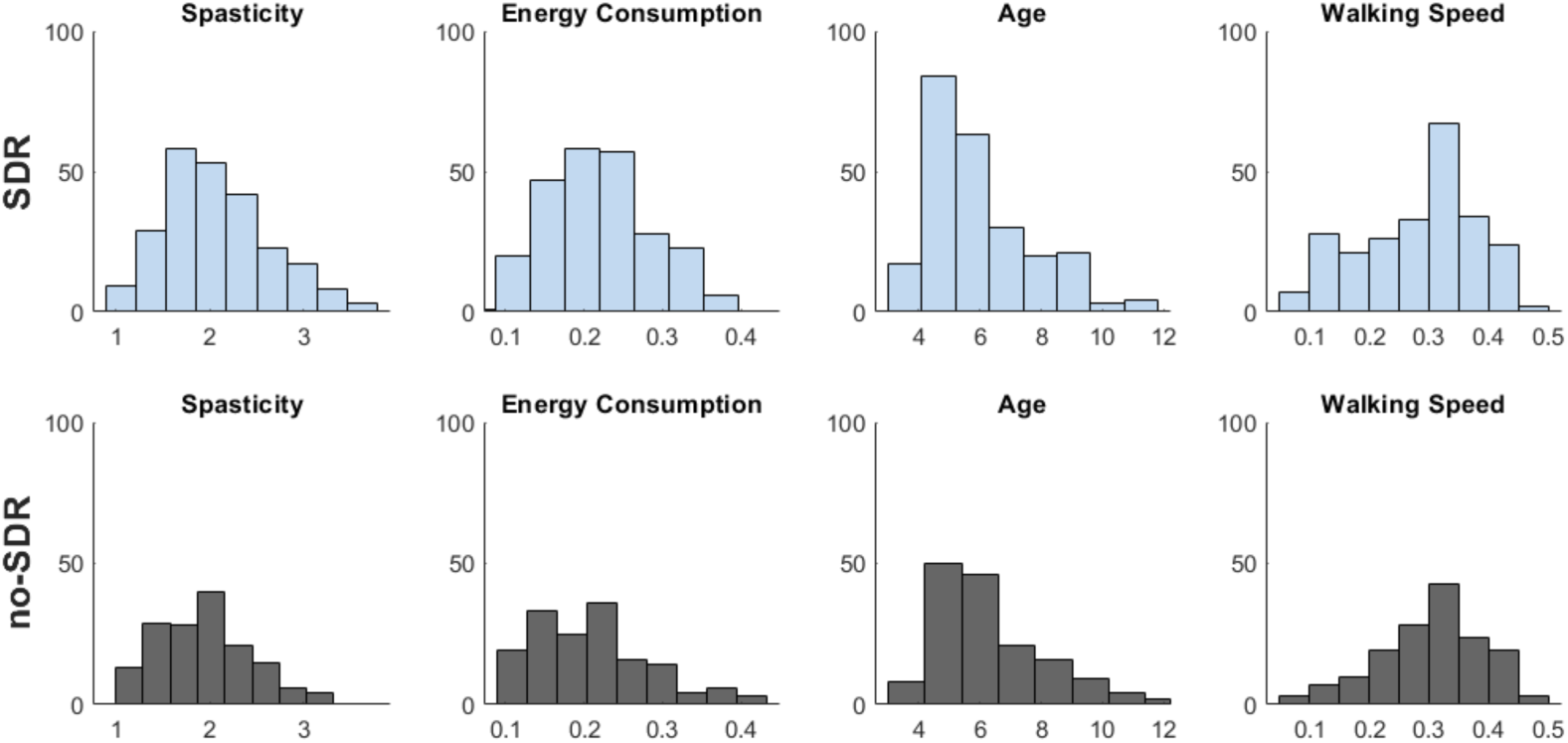
Baseline age (years), net-nondimensionalized (NN) energy consumption, summary spasticity score, and walking speed (m/s) for ***(top)*** children who underwent an SDR and ***(bottom)*** matched peers with CP. Participants were matched for age, NN energy consumption, and summary spasticity score.

### 3.2 Energy Consumption

Energy consumption remained similar between groups at follow-up: 0.18 (SD 0.06) and 0.18 (SD 0.06) for the SDR and no-SDR groups, respectively (Fig. 2). Energy consumption decreased significantly for both groups between visits: SDR, −0.03 (SD 0.07), *P*<0.001, no-SDR, −0.03 (SD 0.06), *P*<0.001. SDR did not result in a greater reduction in energy consumption compared to matched peers (*P*=0.9).

**Figure 2.**
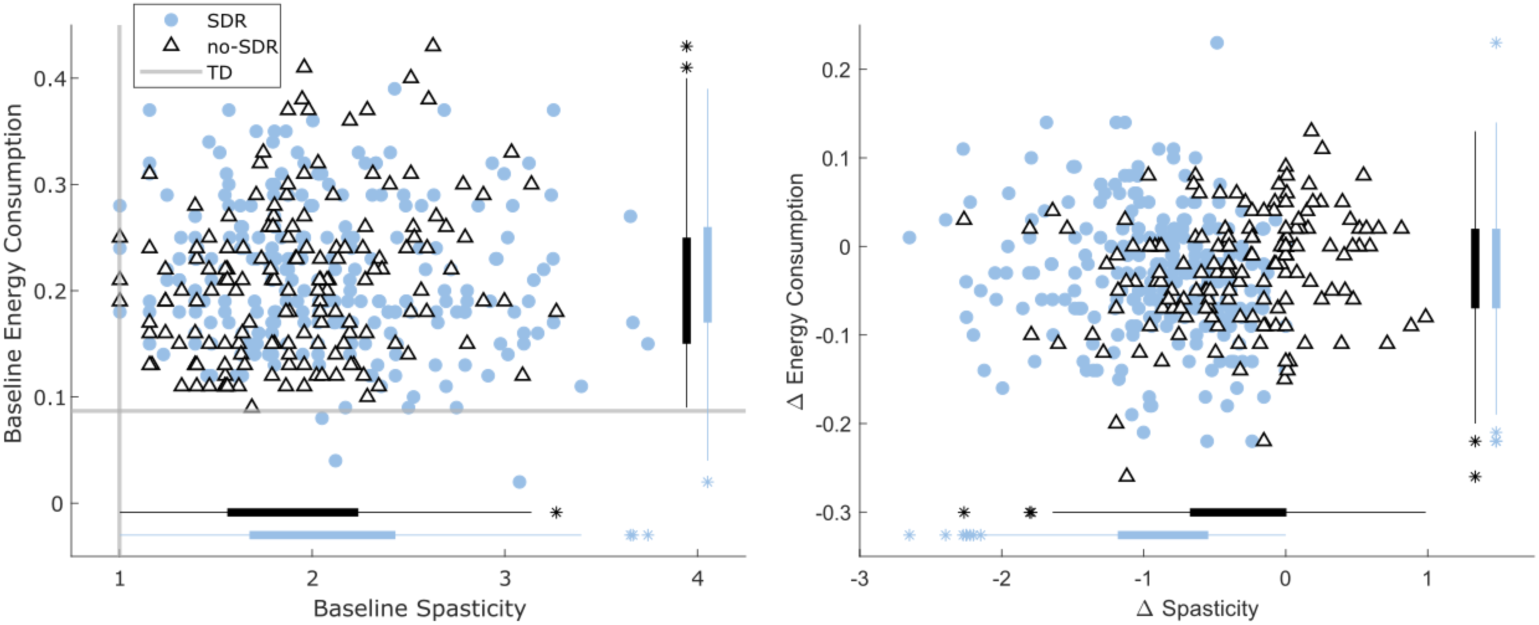
Spasticity and energy consumption for children with CP who underwent an SDR and matched peers with CP. **(*left*)** Baseline spasticity and net-nondimensionalized (NN) power were similar between groups. Gray lines show normative values for typically developing (TD) peers from Gillette. **(*right)*** Spasticity and energy consumption decreased significantly at follow-up for both groups. The SDR cohort had a significantly greater decrease in spasticity compared to the no-SDR group, but a similar decrease in energy consumption. Bars represent distributions for each group including outliers (*).

### 3.3 Spasticity

Children who underwent SDR did have greater reductions in spasticity at follow-up compared to matched peers with CP (*P* < 0.001). The baseline summary spasticity score was 2.1 (SD 0.6) for the SDR group and 1.9 (SD 0.5) for the no-SDR group. The SDR group exhibited a significant decrease in spasticity after surgery with a follow-up summary spasticity score of 1.2 (SD 0.3, *P* < 0.001). The change in spasticity after SDR varied from −2.6 to 0 (change = −0.9 (SD 0.5). The no-SDR group did have a significant decrease in spasticity at follow-up with a summary spasticity score of 1.6 (SD 0.6) (change = −0.3 (SD 0.6), ranging from - 2.3 to +1.0.

### 3.4 Walking Speed

Since walking speed was not a matching variable but can influence energy consumption, we also evaluated changes in walking speed between visits. The baseline nondimensional walking speed was 0.28 (SD 0.09) and 0.30 (SD 0.09) for the SDR and no-SDR groups, respectively. The follow-up nondimensional walking speed was 0.30 (SD 0.10) and 0.30 (SD 0.08) for the SDR and no-SDR groups respectively. There was no significant difference in walking speeds between groups at follow-up (*P*=0.73). While there was a significant increase in walking speed only for the SDR group (*P*=0.001), this increase was small and below the threshold for clinical significance (Oeffinger et al., 2008).

## 4 Discussion

While SDR is effective at reducing spasticity, there was no associated decrease in energy consumption compared to peers with CP who did not undergo an SDR. These results suggest that spasticity is not a primary factor contributing to high energy observed among people with CP. We had hypothesized that if spasticity contributes to energy, then an SDR should result in lower post-treatment energy consumption compared to matched controls with CP. However, the change of energy consumption post-treatment of the children with CP who underwent SDR was not significantly different from the change in energy consumption that the matched controls with CP experienced. This indicates that spasticity is not a primary factor contributing to the increased walking energy in children with CP when compared to TD peers. These findings are consistent with the idea that energy consumption decreases with age among children with CP, independent of treatment (Kamp et al., 2014).

Prior research on the impact of SDR on energy consumption has also reported reductions in energy, but compared to a control group of TD peers or no control group (Carraro et al., 2014; Nicolini-Panisson et al., 2018; Wang et al., 2018). For example, Trost and colleagues (2008) reported a 22% reduction in energetic cost after SDR (Trost et al., 2008). However, the average age before and after SDR was 7.25 (SD 2.1) years and 8.8 (SD 0.4) years, during which we would expect energy to decrease among children with CP. To our knowledge, no studies have included a control group of peers with CP when looking at the impact of SDR on energy consumption. These results demonstrate the critical importance of identifying and comparing to a cohort of peers with CP when evaluating treatments, to differentiate whether changes in function are due to the treatment or natural development. Performing an analysis similar to previous studies with no control group, on the SDR group did show a significant reduction (*P*<0.001) in energy consumption.

There are a few possibilities of why spasticity does not affect energy consumption: a) the additional muscle activity from spasticity isn’t increasing energy consumption, b) the additional muscle activity from spasticity is not large enough to make a large effect, or c) other factors beyond spasticity are the main contributors to the elevated energy consumption in CP. Other possible contributors to elevated energy consumption in children with CP are poor selective motor control, excessive co-contraction, altered muscle properties, or cardiovascular factors. While SDR provided a platform to evaluate the impacts of spasticity, other strategies will be necessary to evaluate the relative importance of other factors and identify optimal strategies for reducing energy consumption for children with CP. This is especially critical, as physical fatigue is very prevalent among individuals with CP throughout their lifespan, and hinders participation and quality of life (Gross et al., 2018; Jahnsen et al., 2003).

This study used retrospective data from clinical gait analysis. Energy consumption is measured in these analyses from a six-minute-walk test that does not control walking speed, mood, or other confounding factors. We matched our groups by baseline spasticity, age, and energy consumption to minimize these effects. Walking speed was not matched for, but there was no significant difference between groups at baseline and follow-up. Further, children walked barefoot which may not represent energy consumption during activities of daily living with braces. However, previous research has shown no significant difference in oxygen consumption between barefoot and shoe (Divert et al., 2008).

## 5 Conclusions

Spasticity was significantly reduced for individuals with CP who underwent an SDR compared to matched peers with CP, but energy consumption was not different between groups. These results demonstrate that spasticity has minimal impact on the high energy consumption observed among children with CP. Both groups demonstrated a reduction in spasticity and energy consumption between visits. Selecting appropriate control groups is critical for research involving children with CP to account for changes in function due to age, development, and other factors. While SDR which is often suggested to reduce spasticity and improve energy, clinicians and families should understand that this procedure does not improve energy consumption during walking.

## 6 Acknowledgments

We thank the staff and clinicians at the James R. Gage Center for Gait and Motion Analysis at Gillette Children’s Specialty Healthcare for their assistance and feedback in preparing this research.

